# Clonal fitness decline in somatic differentiation hierarchies

**DOI:** 10.1101/2023.09.27.559763

**Authors:** Iftikhar Ahmed, David Dingli, Weini Huang, Benjamin Werner

## Abstract

The concept of clonal fitness is fundamental to describe the evolutionary dynamics in somatic tissues. It is now well established that otherwise healthy somatic tissues become increasingly populated by expanding clones with age. However, the dynamic properties and respective fitnesses of these clones are less well understood. Here we show, that in somatic tissues organised as a differentiation hierarchy, theory predicts a natural decline of effective clonal fitness over time in the absence of additional driver events. This decline is intrinsic to the tissue organisation and can be captured quantitatively by a simple heuristic equation that is proportional to 1/time. We also show that the expected fitness decline is directly observable in human haematopoiesis. The predicted short and long term dynamics agree with *in vivo* observations using data of Neutrophil recovery after bone marrow transplants and naturally progressing Chronic Lymphocyte Leukemia (CLL). We further show that theory predicts the existence of a long term equilibrium fitness. All CLL patients transition into a stable equilibrium fitness eventually. We find significant inter-patient variation of long term fitness and a strong correlation with disease aggressiveness. Interestingly, CLL long term fitness can be forecast based on the early stages of disease progression, suggesting a Big Bang like model for CLL evolution.

## Introduction

The expansion of clones in somatic tissues with age is a well-documented phenomenon and universal to all tissues [1–3]. It is a naturally expected outcome of somatic evolutionary processes, underlying tissue transformation, and cancer progression [4–6]. However, although clonal expansions are common,cancers are overall still rare, suggesting most clonal expansions do not progress [7]. This is an interesting evolutionary phenomenon and the containment of clones has to be understood in the context of its somatic environment [8].

Most human tissues are hierarchically organized [9–11]. A comparably small population of stem cells maintains tissue homeostasis through a balance of self-renewal and differentiation, giving rise to fully functional differentiated cells [12]. Such structures minimize the accumulation of mutations while at the same time allowing the production of enormously large numbers of fully differentiated cells within a short time [13–16]. This is possible because only mutations in stem cells persist [17]. All non-stem cell derived mutations vanish in the long run as the lifetime of any non-stem cell is finite (unless the cells acquire stem cell like properties due to mutations). While clonal dynamics have to be understood in the context of such differentiation hierarchies, we focus on the clonal dynamics within the hematopoietic system. There are three general patterns of clonal trajectories naturally occurring in such hierarchies, continued clonal expansion, clonal homeostasis, and waves of clonal extinction [11].

We investigate how a differentiation hierarchy affects clonal fitness. We show that mathematical models predict a continuous decline of clonal fitness with age in a differentiation hierarchy in the absence of additional driver events. This decline seems universal and is expected to occur even for aggressively exponentially expanding clones. In the long run, clonal dynamics approach a constant equilibrium fitness that is either positive for continued clonal growth or negative for waves of clonal extinctions.

We then use data from two different patient cohorts to test our theoretical expectations directly in human hematopoiesis. We first follow neutrophil count recovery after stem cell transplantation in 19 patients with multiple myeloma who received high dose melphalan conditioning followed by the infusion of autologous stem and progenitor cells. These data represent the earliest stages of a clonal expansion and we observe a sharp decline in clonal fitness in all patients. To test the predictions of a long-term equilibrium of clonal expansions, we analyze clonal fitness in 85 patients of chronic lymphocytic leukemia (CLL), some of whom remained untreated for up to 20 years. All patients transitioned into a stable equilibrium fitness. Furthermore, equilibrium fitness is also correlated with disease aggressiveness, which allows to forecast of disease progression early.

## Results

### Clonal fitness in hierarchical tissues

The dynamics of cells in hierarchically organized tissues can be modeled mathematically by compartmentalized structures. Each compartment represents cells at certain stages of differentiation. Cells move at predefined rates between compartments. Usually, homeostasis in such hierarchies is maintained by a self-renewing population of stem cells that give rise to all fully differentiated cells within a tissue. The deterministic dynamics of cells in such hierarchies can then be captured by a set of differential equations [11], given by

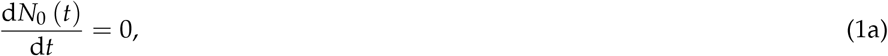

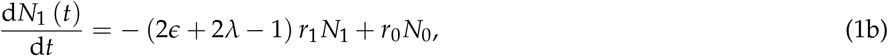

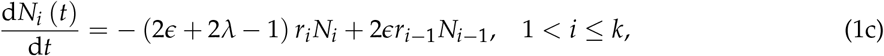

Here, *N*_*i*_(*t*) denotes the number of cells in compartment *i* at time *t* (0 ≤ *i* ≤ *k*), *ϵ* is the cell differentiation probability, *λ* is the cell death probability, and *r*_*j*_ is the cells proliferation rate in compartment *j*. Thus, the probability of self-renewing is (1 − *ϵ* − *λ*). We introduce the notation *α* for the effective net growth rate per compartment as *α* = *ϵ* + *λ* − (1 − *ϵ* − *λ*) = 2*ϵ* + 2*λ* − 1.

We are interested in the dynamics of clones within such differentiation hierarchies. If one places a cell with certain proliferation parameters at any stage within this hierarchy, what are the dynamics of its progeny? Suppose *N*_*ij*_ is the size of that clonal population in compartment *i* when the first cell of that clone originated in compartment *j*. Formally this corresponds to the initial conditions

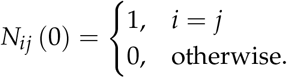

Together with equations (1) this leads to a solution of the form

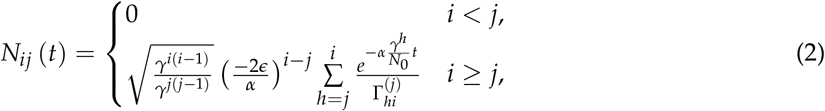

where we set 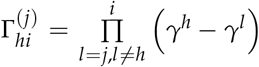 with *r*_*i*_ = *r*_0_*γ*^*i*^ in equation (1c). The set of equations (2) allows for three different clonal trajectories that depend on the value of the differentiation rate *ϵ*. For *ϵ* < 0.5, cell numbers are increasing exponentially across compartments. For *ϵ* = 0.5, we have homeostasis where the cell number reaches a constant equilibrium. Wherever *ϵ* > 0.5, we observe waves of clonal extinction traveling through compartments, where clonal waves result from a lack of sufficient self-renewal, causing clonal extinction in the long run, see Figure 2a for an example.

It is then natural to ask, if we have *in vivo* time series data and want to estimate the fitness of clonal expansions with potentially important prognostic consequences for patients, what should we expect? Theory predicts that the experimentally observed fitness will critically depend on the exact timing of the experiment despite the underlying intrinsic parameters of the clonal expansion remain unchanged.

To show this formally, we define the fitness *s* of cells in a differentiation hierarchy as the change of cell numbers within a time interval ∆*t* normalized by its mean population size

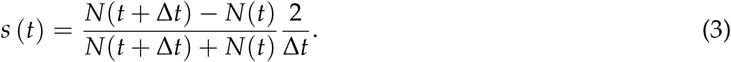

A value of *s* = 0 corresponds to homeostasis, *s* > 0 to growing clonal expansions and *s* < 0 to shrinking clonal populations. In the limit of ∆*t* → 0 and normalizing for population size *N*, this fitness becomes the per capita growth rate of cells and is given by

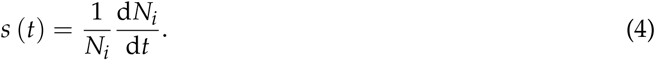

Using equation (4) and considering any intermediate compartment *i* in equations (1), we have

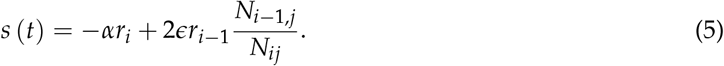

Substituting equation (2) and simplifying we obtain

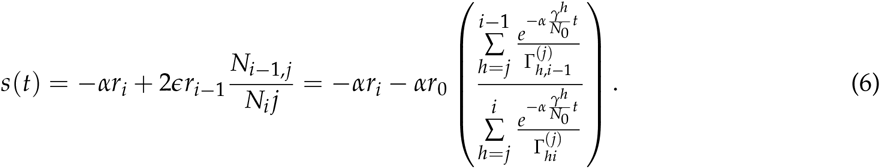

We show two representative examples of the dynamic behavior of equation (6) in Figure 2b.

Although all intrinsic parameters of the clonal population, e.g. the proliferation rate and differentiation probability, are kept constant across all compartments, fitness is predicted to decline over time in the absence of additional events. This decline is universal. It occurs for exponentially expanding populations as well as waves of clonal extinctions (Figure 2a & 2b). Furthermore, the decline decelerates with time. For sufficiently long times, equation (6) predicts the existence of two distinct equilibria that are clonal expansions (*ϵ* < 0.5) or waves of clonal extinction (*ϵ* > 0.5) within a differentiation hierarchy ultimately reach

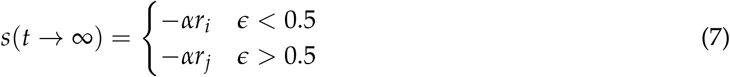

Equations (7) show two important differences for the long term fitness of clonal expansions and extinctions. Firstly, the fitness of clonal expansions always remains positive, whereas the long-term fitness of waves of clonal extinction is negative. Secondly, clonal expansions are ultimately dominated by the proliferation of the most differentiated cells *r*_*i*_, whereas clonal extinctions depend on the proliferation of the slowest least differentiated cells *r*_*j*_.

Although being exact, the change of the clonal fitness function, equation (6), is rather involved and in principle depends on all microscopic parameters, which makes direct comparisons to time series data with unknown microscopic parameters difficult. Heuristically, equation (6) can be approximated by

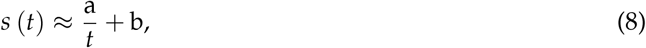

with only two free parameters, a and b (Figures 1b & 2c). The first parameter a corresponds to the effective rate of fitness decline and the second parameter b represents the equilibrium fitness of clonal dynamics. This heuristic approximation captures the dynamics of the clonal fitness well and allows us to numerically fit time series data of clonal dynamics to derive estimates for the rate of decline a and the long term fitness b, Figure 2c & 2d.

**Figure 1.**
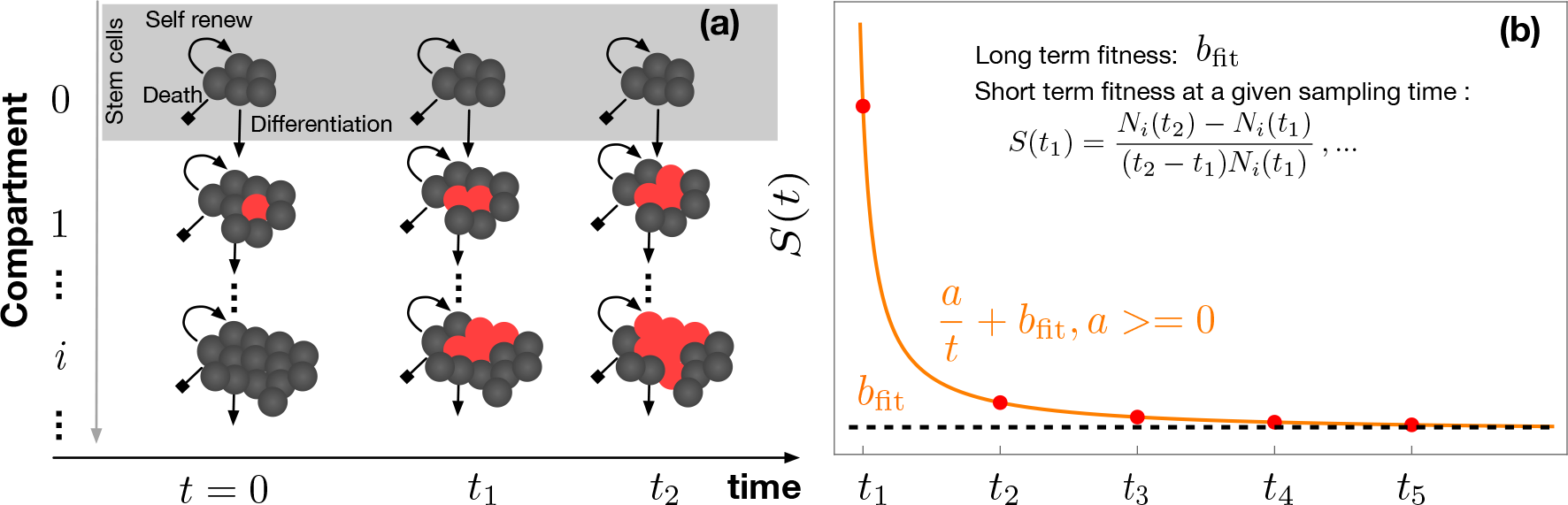
Clonal expansions and differential fitness in hierarchically organised tissues. Conceptual figure showing **(a)** clonal expansions in a differentiation hierarchy and **(b)** the concept and dynamics of the differential fitness of these expanding clones. Theory predicts a decline of clonal fitness over time that is proportional to 1/time and can be heuristically described by the simple equation 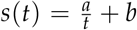 with a long term equilibrium clonal fitness b.

**Figure 2.**
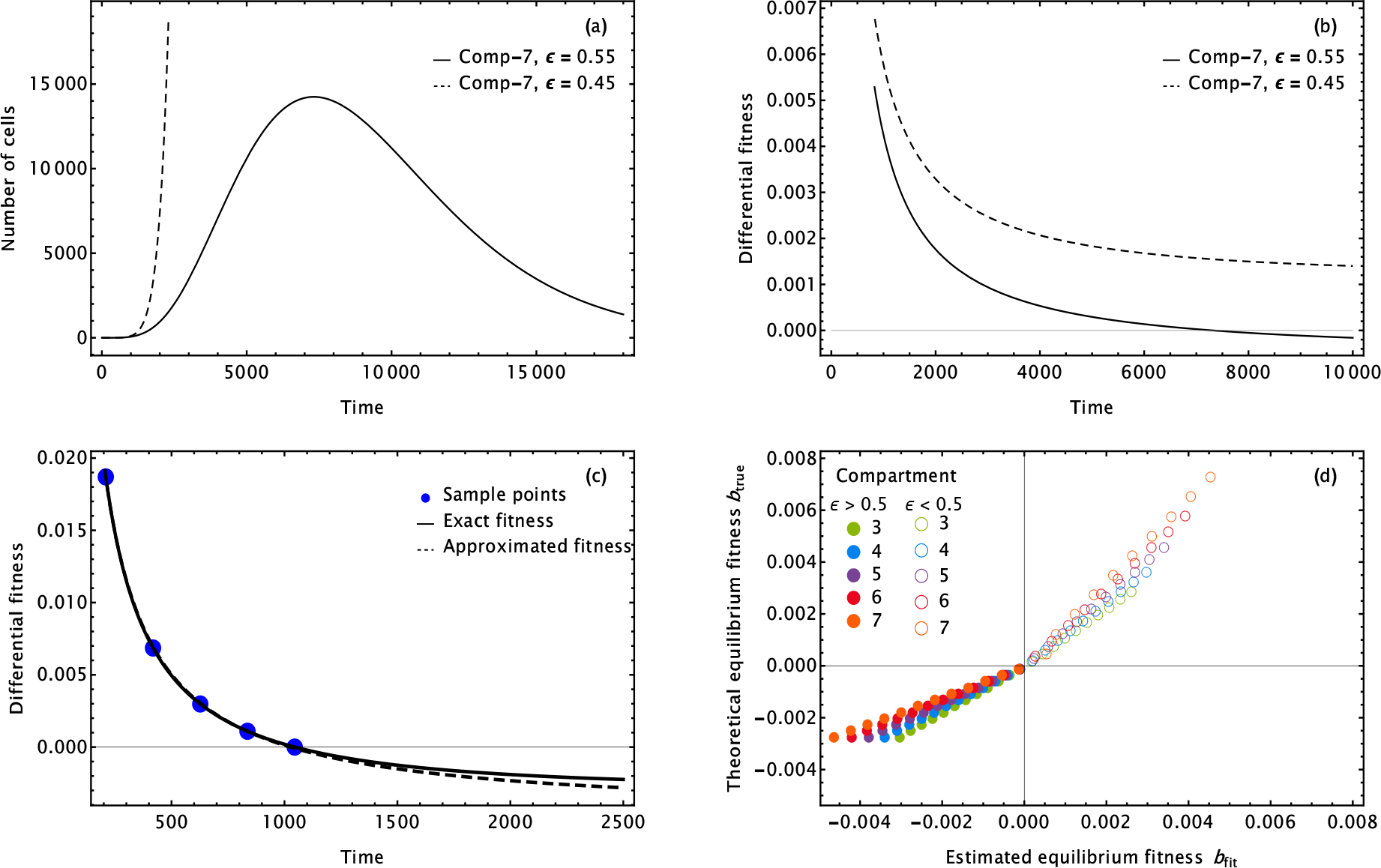
Representative realisations of theoretical predictions of clonal expansions and differential fitnesses in hierarchical tissues. **(a)** Realisations of eq. (2) in compartment 7. Dashed line represents an exponentially growing clone (*ϵ* = 0.45) and the line shows a wave of clonal extinction *ϵ* = 0.85. **(b)** Realisation of differential fitness (eq. (4)) for *ϵ* = 0.55 (black line), and *ϵ* = 0.45 (dashed line). **(c)** Clonal fitness decline is well described by the heuristic approximation *s*(*t*)= *a/t* + *b*. **(d)** Long term equilibrium fitness can be forecast using data on the early expansion of clones.

Equation (8) also allows us to estimate the time *τ* till equilibrium fitness is reached. We say that we are in equilibrium if the change in clonal fitness is below a certain threshold *h*. We then have

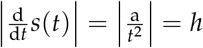 and thus the time to equilibrium *τ*h is given by

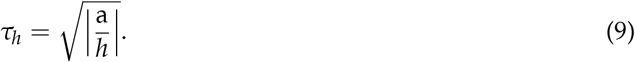

Below, we will use *h* = 0.01 as the threshold for the equilibrium phase.

### Neutrophil recovery after stem cell transplantation

Our theoretical modelling suggests that the fitness of clonal expansions declines sharply early and levels off towards the equilibrium value for sufficiently long times. To test the prediction of a sharp initial fitness decline of clonal expansions in human haematopoiesis, we measured the neutrophil counts in 19 patients with multiple myeloma who received autologous stem and progenitor cells (CD34+) after myeloablative conditioning with melphalan. This is standard therapy for fit patients with multiple myeloma where patients initially receive induction therapy and following disease control they have autologous stem and progenitor cells collected via growth factor mobilization (e.g. granulocyte colony stimulating factor). Typically 3 × 10^6^CD34 cells are required for a single stem cell transplant. Initially, the neutrophil (and other blood cell counts) fall due to myeloablation but then typically around 13 days after the stem cell infusion, neutrophil recovery is observed. Thus patients are monitored carefully for their neutrophil recovery during this period. For a number of days, the neutrophil count will be too low for accurate quantitation. This is known as the neutrophil nadir. Data from 19 such patients was captured for this analysis.

The recovery of neutrophils after depleting all the bone marrow resembles a clonal expansion in the background of an empty differentiation hierarchy. Neutrophils were measured daily within the first 3 weeks after the stem cell transplant, allowing us to use equation (4) to calculate the change of clonal fitness over time across patients. Time trajectories were fitted using the heuristic approximation (8) to each patient separately. All patients received their stem cell infusion on day 0 and neutrophil counts of the re-expanding transplant usually remain below the detection threshold for the first 10 days. After day 10, neutrophil counts are detectable and rise steadily. As predicted by the modelling, the fitness of these early clonal expansions is decreasing rapidly in all but two patients, Figure 3a & 3b (see also Figure S2 for a summary of all patients). Typically neutrophil fitness becomes negative at day 14 to 15 after the autologous stem cell transplant. However, the time to equilibrium is considerably longer 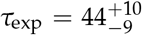 (days) with considerable variation between patients (0 to 78 days), Figure 3c. This is in keeping with clinical observations that patients may take several weeks for their neutrophil count to return to normal and stay there.

**Figure 3.**
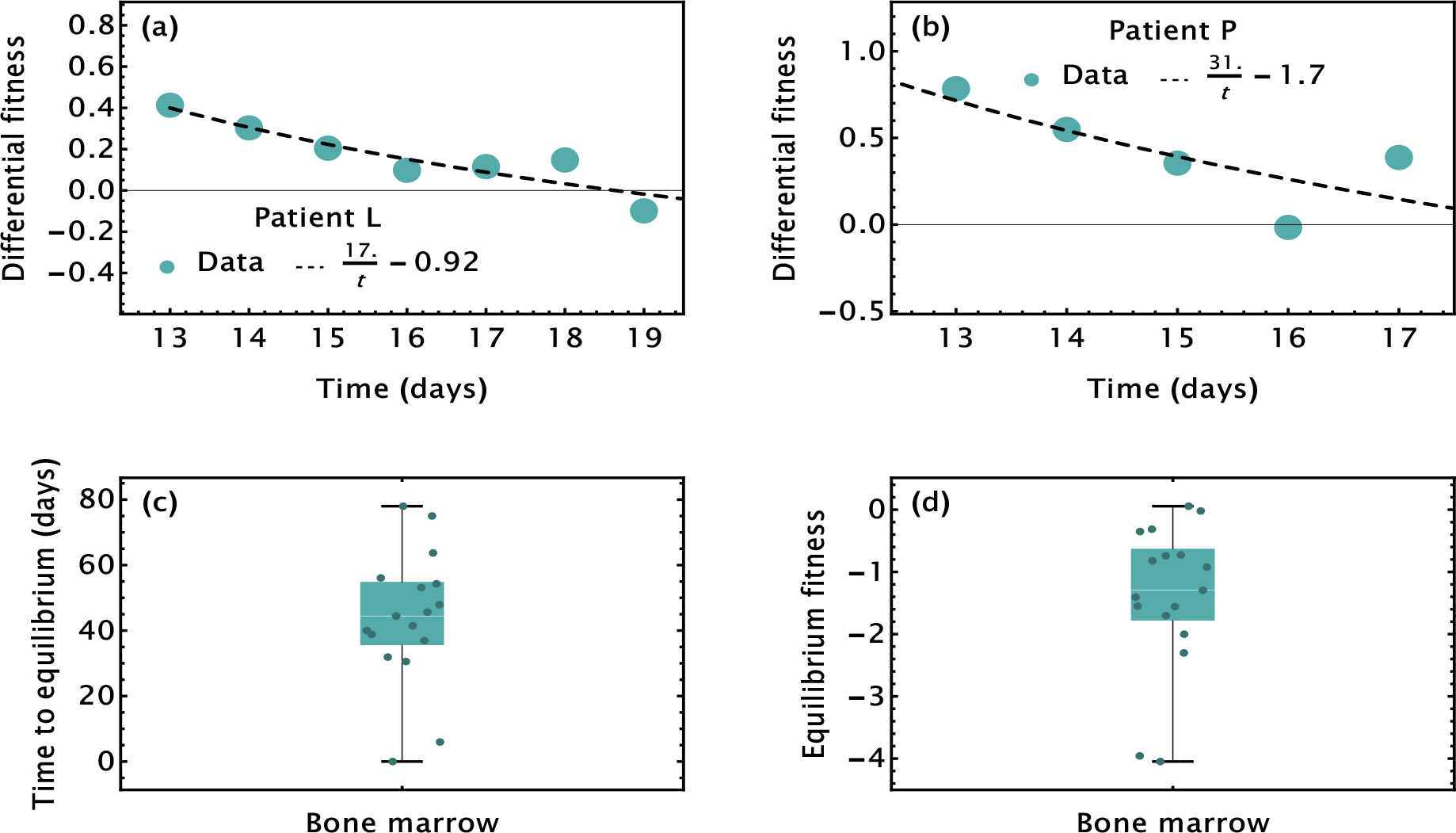
Neutrophil recovery after bone marrow transplant. **a),b)** Two examples of typical differential fitness trajectories of Neutrophil recovery after stem cell transplant. Differential fitness is following the heuristic 1/t decline and drops sharply within a few days. **c)** Time to equilibrium is estimated to be 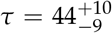 days. **d)** Long term clonal fitness of Neutrophil recovery becomes negative 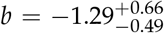 per day.

### Equilibrium clonal fitness of Chronic Lymphocytic Leukemia

To test the prediction of the long-term equilibrium fitness of clonal expansions in a differentiation hierarchy, we analyzed the trajectories of 85 naturally progressing patients with chronic lymphocytic leukemia (CLL) originally published in [18]. CLL patients are often monitored for many years without intervention, enabling us to observe the long-term change in clonal fitness retrospectively. White blood cell counts in this cohort were followed for up to 20 years with often multiple measurements per year. In the original study, the patient cohort had been retrospectively separated into three categories, exponential growth, logistic growth, and indeterminate growth [18]. These three categories correlated with the aggressiveness of the disease. Patients exhibiting exponential growth patterns were more likely to require treatment compared to patients with logistic clonal growth. Exponential growth also correlated with a higher number of detectable driver mutations compared to logistically progressing disease [18]. Patients with indeterminate growth patterns had significantly shorter follow-up times, see Supplementary Figures S(11 – 14). Thus, this category likely contains both logistic and exponential growth patterns that were not yet distinguishable in the original study. In our subsequent analysis, we use the same grouping of patients by the three growth categories as in the original study.

As predicted by our theory, a constant clonal equilibrium fitness is reached in all patients independent of classification. Summaries of all clonal fitness over time are shown in Supplementary Figures S(3 – 14). The initial fast decline of the clonal fitness is less pronounced in most CLL patients compared to the clonal fitness of neutrophils after bone marrow transplants. This is likely due to the normal physiologic expansion of neutrophils after transplantation compared to the oncogene driven growth of the cells in CLL. The sharp initial drop is most pronounced at the earliest stages of clonal expansions. In most CLL patients, the diseases had been present for some time before they entered monitoring and data collection.

We then ask if differences in disease aggressiveness are also reflected in the long-term fitness of clonal expansions. By fitting equation (8) to individual fitness trajectories, we estimate the long-term fitness b for all 85 patients. A summary of this fitness is shown in Figure 4. The equilibrium fitness of logistically and exponentially growing clones differ significantly (*p* = 2 × 10^−5^, T-test) Figure 4c. The median clonal fitness of exponential clones is 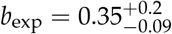 (per year), whereas for logistic clones we find 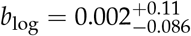 (per year). The clonal fitness for exponentially growing tumours corresponds to an approximate exponential growth rate of the disease, allowing us to estimate doubling times, which are in the range of 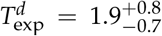 (years), which strongly correlate with adverse outcome [19, 20]. Furthermore, clonal fitness is always positive in patients exhibiting exponential growth, whereas for logistically growing clones, fitness can be negative or positive, and between patients, variation is higher (Figure 4c). However, most fitness in the logistic growth category cluster around b = 0, suggesting either slow expansions if *b*_log_ is small and positive, or an approximately constant disease if *b*_log_ is small and negative. There are 3 patients in the logistic category with *b*_log_ < − 0.2 (per year), suggesting possible clonal extinctions of CLL in these patients in the long-term. However, the extinction time is on the scale of decades and residual disease might remain detectable throughout life. The clonal fitness of patients with indeterminate growth patterns is between logistic and exponential growth 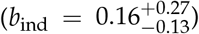 and is not significantly different from either logistic or exponential growth (Supplementary Figure S16a). This is in line with the idea that the indeterminate category is a mix of patients with logistic or exponential growth patterns respectively because of the significantly shorter follow-up times.

**Figure 4.**
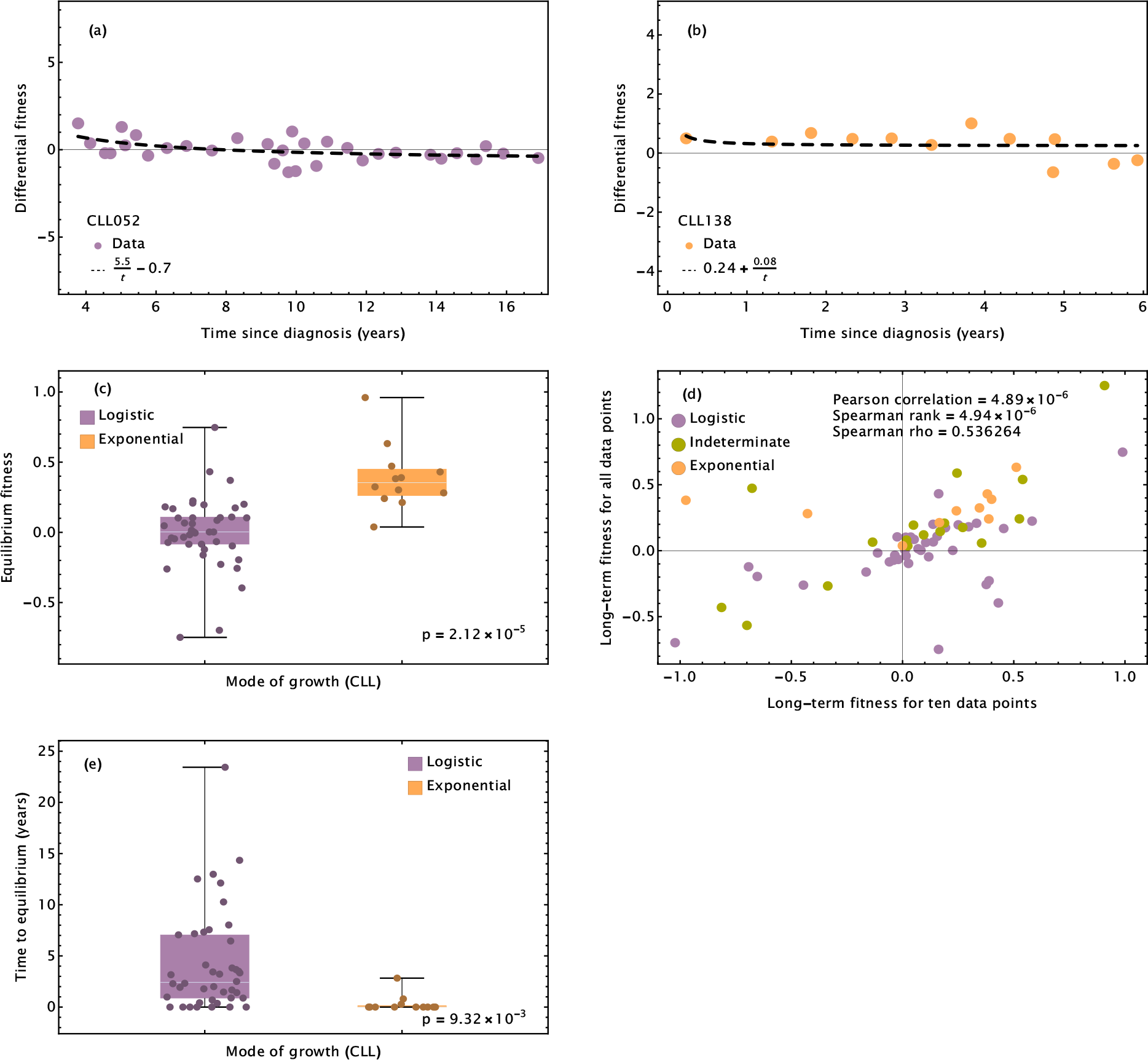
Clonal fitness in CLL. Fitness trajectory of a **a)** logistically and **b)** exponentially expanding CLL. Trajectories of all 85 patients are shown in Supplementary Figures S(3 –14). **c)** Equilibrium fitnesses of exponentially progressing CLLs are always positive (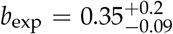 per year) and significantly higher than logistically progressing CLL (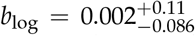 per year),(*p* = 2 × 10^−5^ T-test).**d)** Equilibrium fitness inferences from early disease stage (first 10 data points) correlate strongly with retrospective fitness estimates (SpearmanRho = 0.54, *p* = 1.4 × 10^−6^). **e)** Time to equilibrium is significantly faster for exponential growth (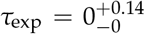 years), compared to logistically progressing CLL(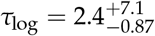 yearsp = 9 × 10−^3^,T-test).

### Clonal fitness forecast in CLL

Having shown that the equilibrium clonal fitness strongly correlates with retrospective grouping of patients disease trajectories, we ask if we can identify exponential or logistic growth early. More precisely, we ask, can the change in clonal fitness at early time points forecast long-term clonal fitness within patients?

We showed that within our mathematical model, such a forecast is possible (Figures 1b and 2d). If we fit heuristic equation (8) to the early dynamics of either exponentially growing clones or waves of clonal extinctions with known model parameters, we can compare estimated *b*_fit_ to the exact *b*_true_, given by equation (6). These estimates correlate well with the theoretical prediction (Spearman Rho = 0.993). We tend to slightly underestimate the rate of clonal expansions and slightly overestimate the rate of clonal extinctions. However, the estimates conserve signs, allowing us to distinguish exponential expansions and clonal extinctions in principle.

We now test if we can forecast disease progression similarly in CLL patients. We use the first, 5, 7, and 10 data points available for each patient and estimate the equilibrium fitness by fitting equation (8) to those early data points, Supplementary Figures S(4 – 6), S(8 – 10), & S(12 – 14). The resulting estimates of the equilibrium fitness b can then be compared to the estimated long-term fitness derived from complete disease trajectories. All forecasts correlate with the corresponding clonal fitnesses from complete disease trajectories (Figures 4d and S15). Using the first 10 data points, we find a strong positive correlation (Spearman Rho = 0.56, *p* = 1.4 × 10^−6^). Even when using only the first 5 data points for each patient, the correlation remains positive and significant (Spearman Rho = 0.43, *p* = 5 × 10^−4^). The strong positive correlation of early and late fitness estimates as well as the existence of a stable equilibrium fitness in virtually all patients suggest that the average fitness CLL is determined early and maybe surprisingly, does not change significantly throughout disease progression. This agrees with very recent observations that even the transition into Richter syndrome, the transformation of CLL into an aggressive diffuse B-cell lymphoma is already encoded early in the evolutionary history of CLL patients [21].

## Discussion

The concept of clonal fitness is fundamental to evolutionary dynamics. Yet, direct measures of fitness remain difficult in biological systems [22]. In somatic evolution, clones with increased fitness populate all aging tissues [1, 3, 23]. A quantitative understanding of these fitness changes is complicated by the fact that somatic tissues are highly structured and regulated. Most clones, even with high initial fitness usually do not expand beyond certain limits. It is thus natural to expect that any observable clonal fitness is not a constant numerical value, assigned to a specific clone *a priori*, but a dynamic quantity that changes as a clone is expanding within its changing environment.

Here, we have shown that differentiation hierarchies naturally reduce clonal fitness over time. This decline is universal and can be observed both during the expansion of healthy Neutrophil populations after stem cell transplant and transformed (malignant) CLL clones. The fitness decline follows a simple heuristic time dynamics that is proportional to 1/time, with only two free effective parameters, the initial fitness decline and the long term equilibrium fitness. Estimating these parameters in 85 patients with naturally progressing CLL, we found that long term fitness correlates strongly with disease aggressiveness and more importantly, long term fitness can be forecast early. This strongly suggests that in the case of CLL, clonal fitness is determined early and does not change much during disease progression. This is in line with previous observations that the acquisition and expansion of additional subclonal driver mutations in CLL is rare [18] and even the transition into Richter syndrome, the transformation of CLL into an aggressive diffuse B-cell lymphoma is already encoded early in the evolutionary history of CLL [21]. This has important implications for monitoring and possibly stratifying patients into different risk and treatment groups by ideally combining genomic information and longitudinal measures of clonal expansion rates over time.

Whether this Big Bang like dynamics [24] and its implications are unique to CLL or are more prevalent across other liquid and solid tumours remains an open question.

## Acknowledgments

I.A. acknowledges support by HEC Pakistan through the project Establishment of University of Swat Phase-I. B.W. is supported by a Barts Charity Lectureship (grant no. MGU045) and a UKRI Future Leaders Fellowship (grant no. MR/V02342X/1).

We are grateful to Catherine Wu and Gad Getz for providing us access to the CLL patient cohort.

## Supplementary Figures

**Figure S1.**
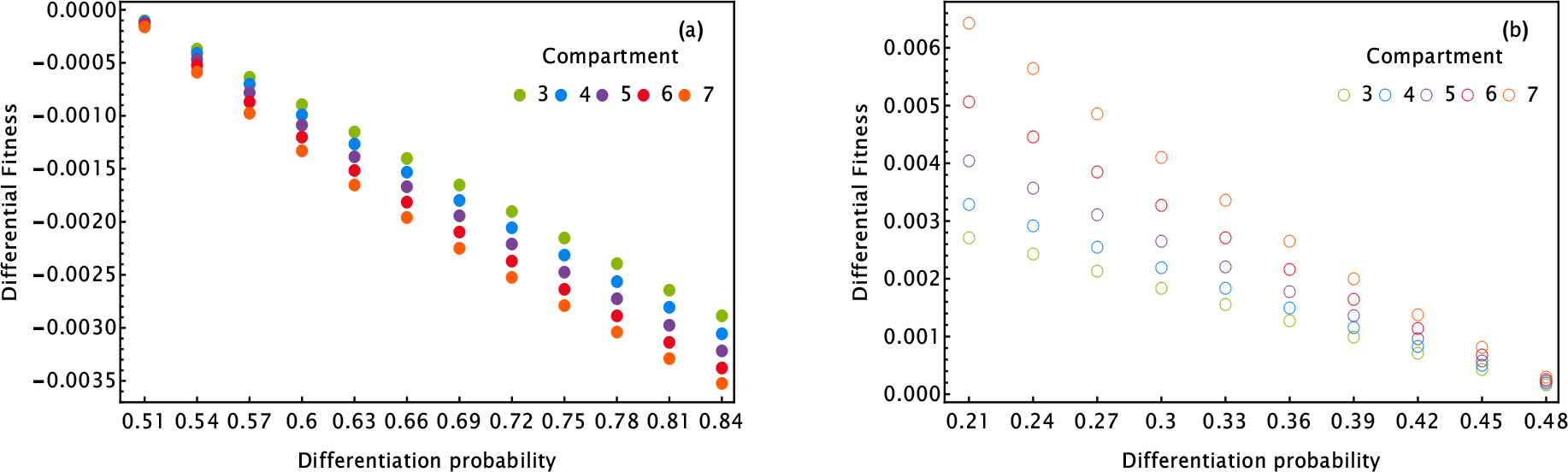
Fitted values of long term fitness in fixed compartment for **(a)** *ϵ* > 0.5 and **(b)** *ϵ* < 0.5.

### Neutrophil fitting all patients

**Figure S2.**
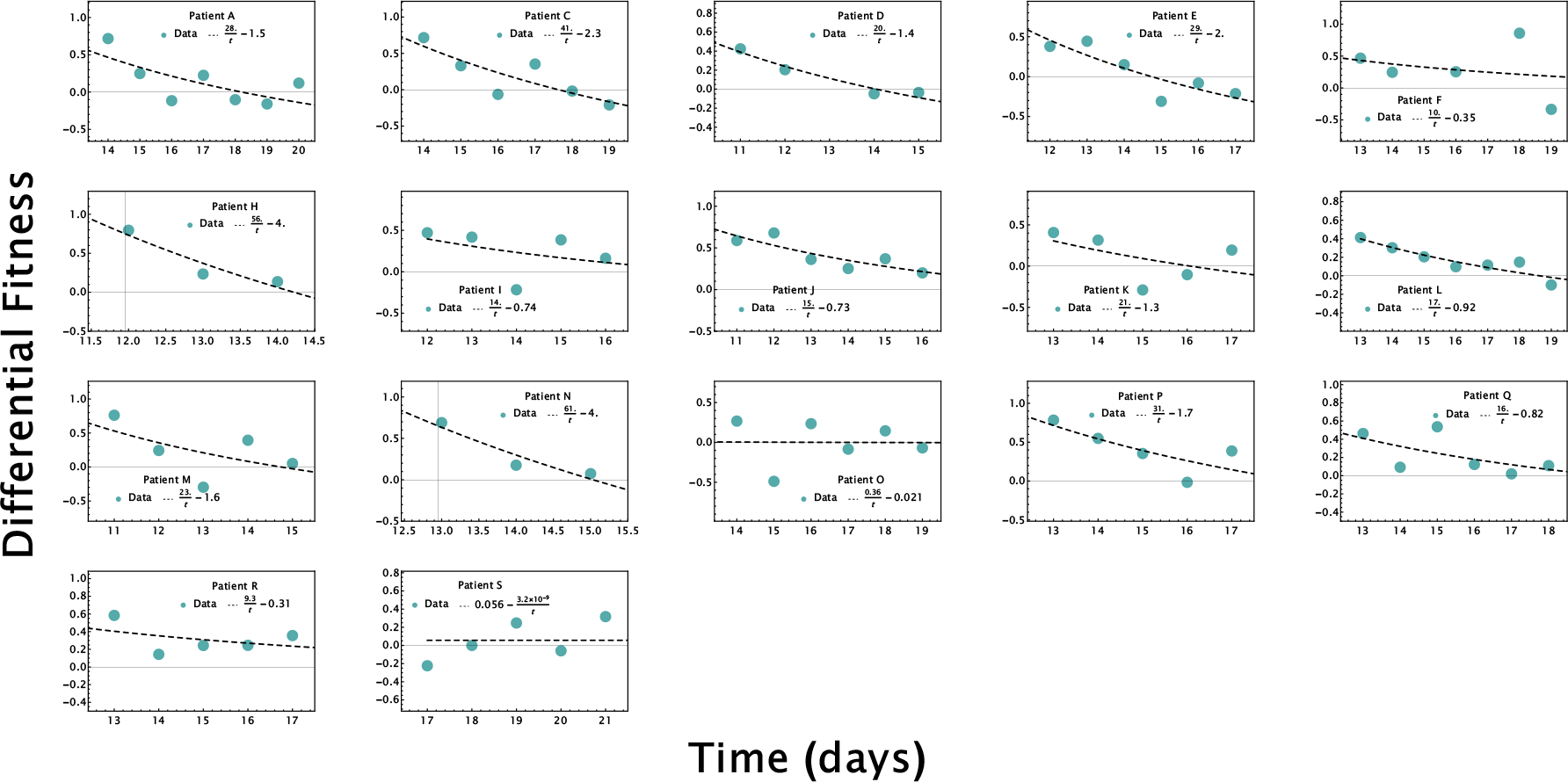
Neutrophil differential fitness following a bone marrow transplant.

### CLL logistic growth pattern

**All points fitting**

**Figure S3.**
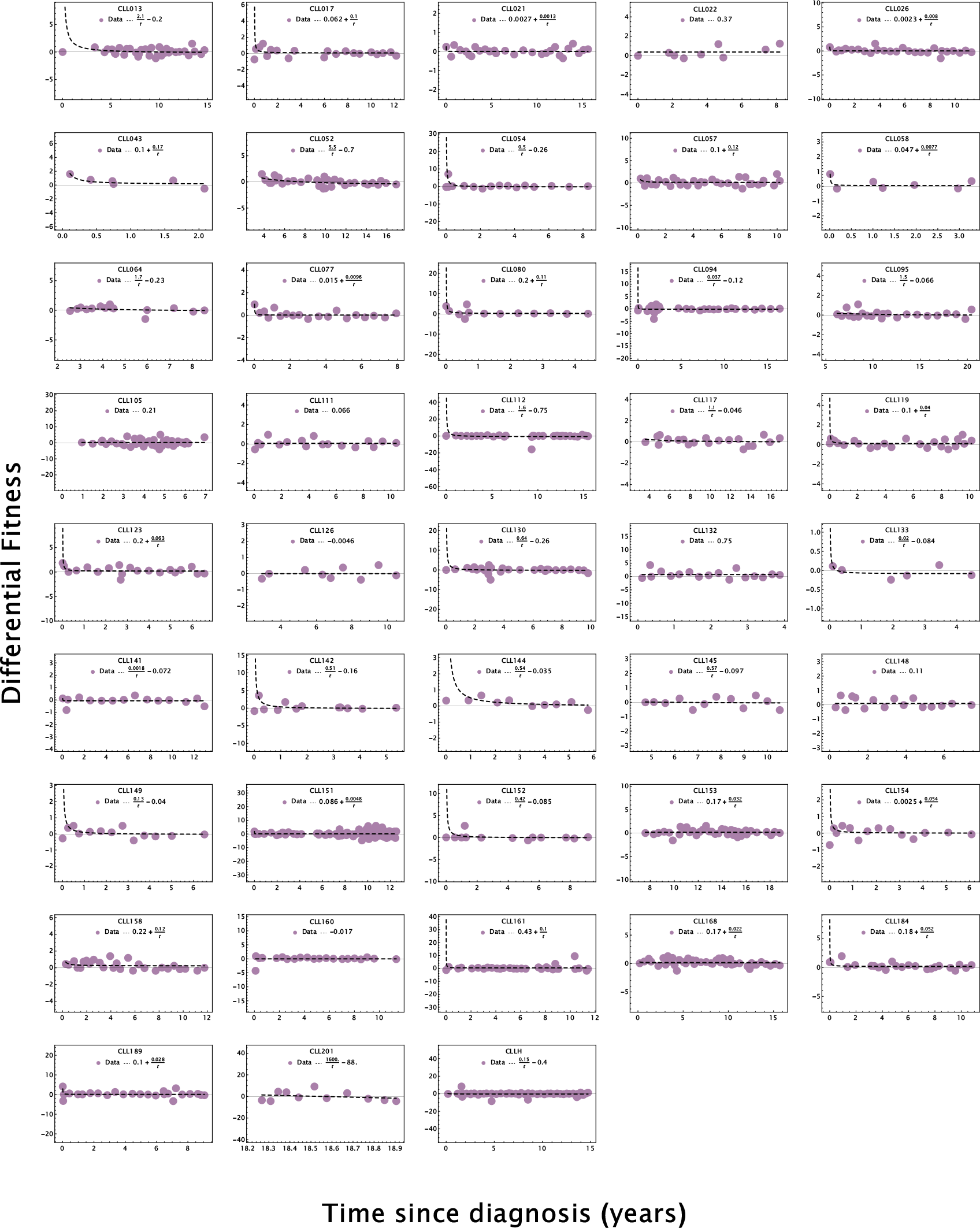
CCL patients with logistic growth pattern. The fitting is based on all available clinical observations

### First 5−points fitting

**Figure S4.**
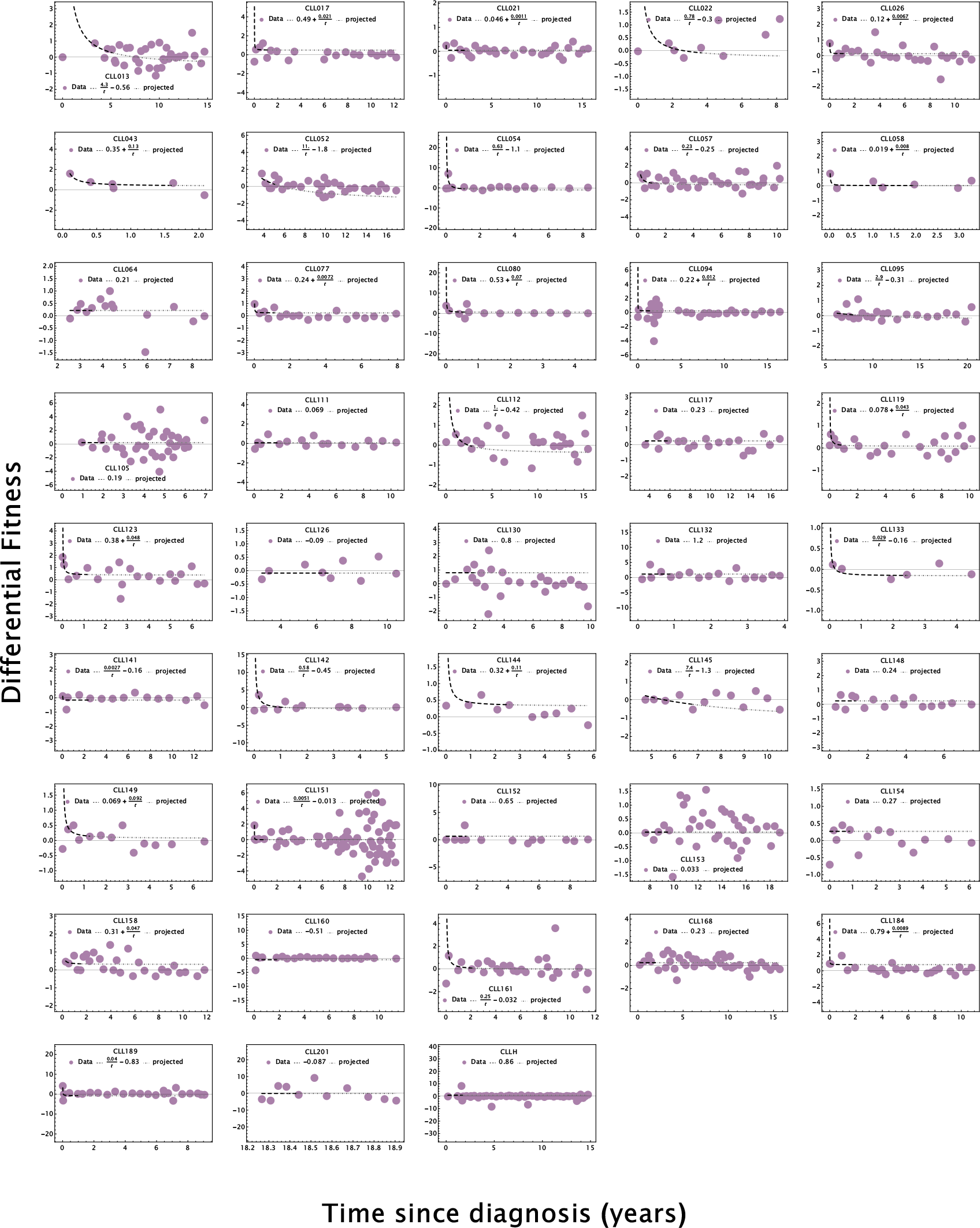
CCL patients with logistic growth pattern. The fitting is based on the first 5 clinical observations.

### First 7−points fitting

**Figure S5.**
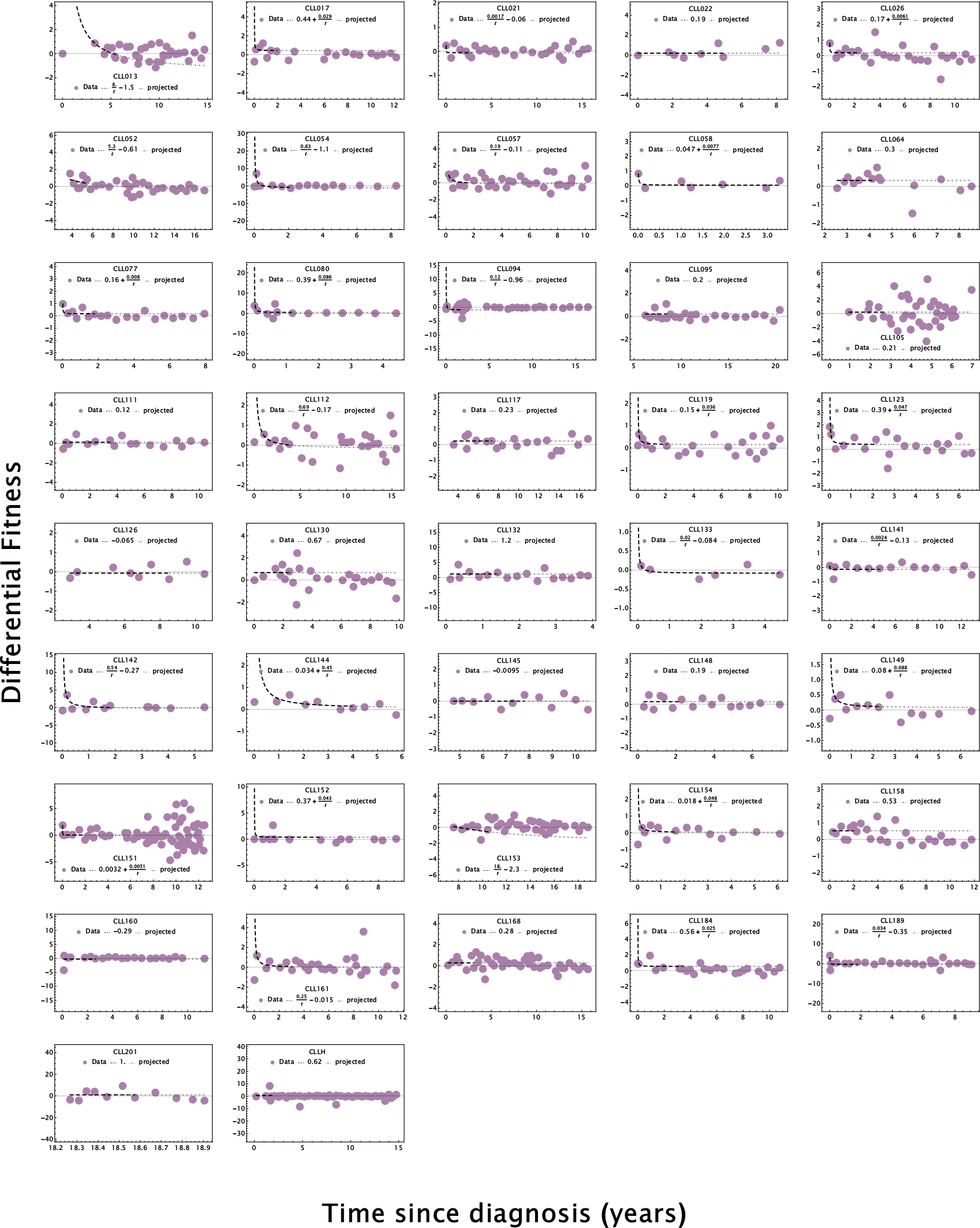
CCL patients with logistic growth pattern. The fitting is based on the first 7 clinical observations.

### First 10−points fitting

**Figure S6.**
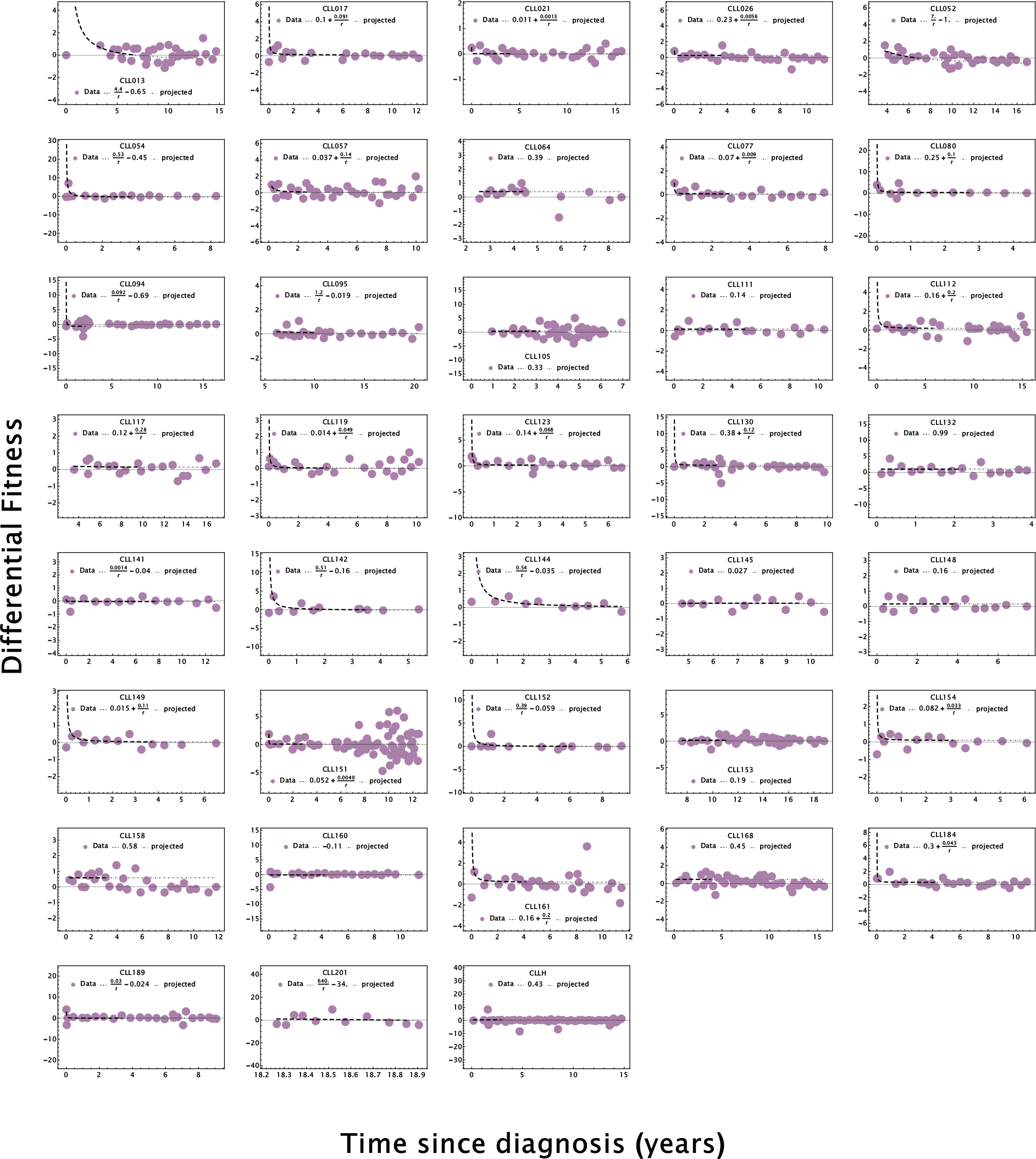
CCL patients with logistic growth pattern. The fitting is based on the first 10 clinical observations.

### CLL indeterminate growth pattern

All points fitting

**Figure S7.**
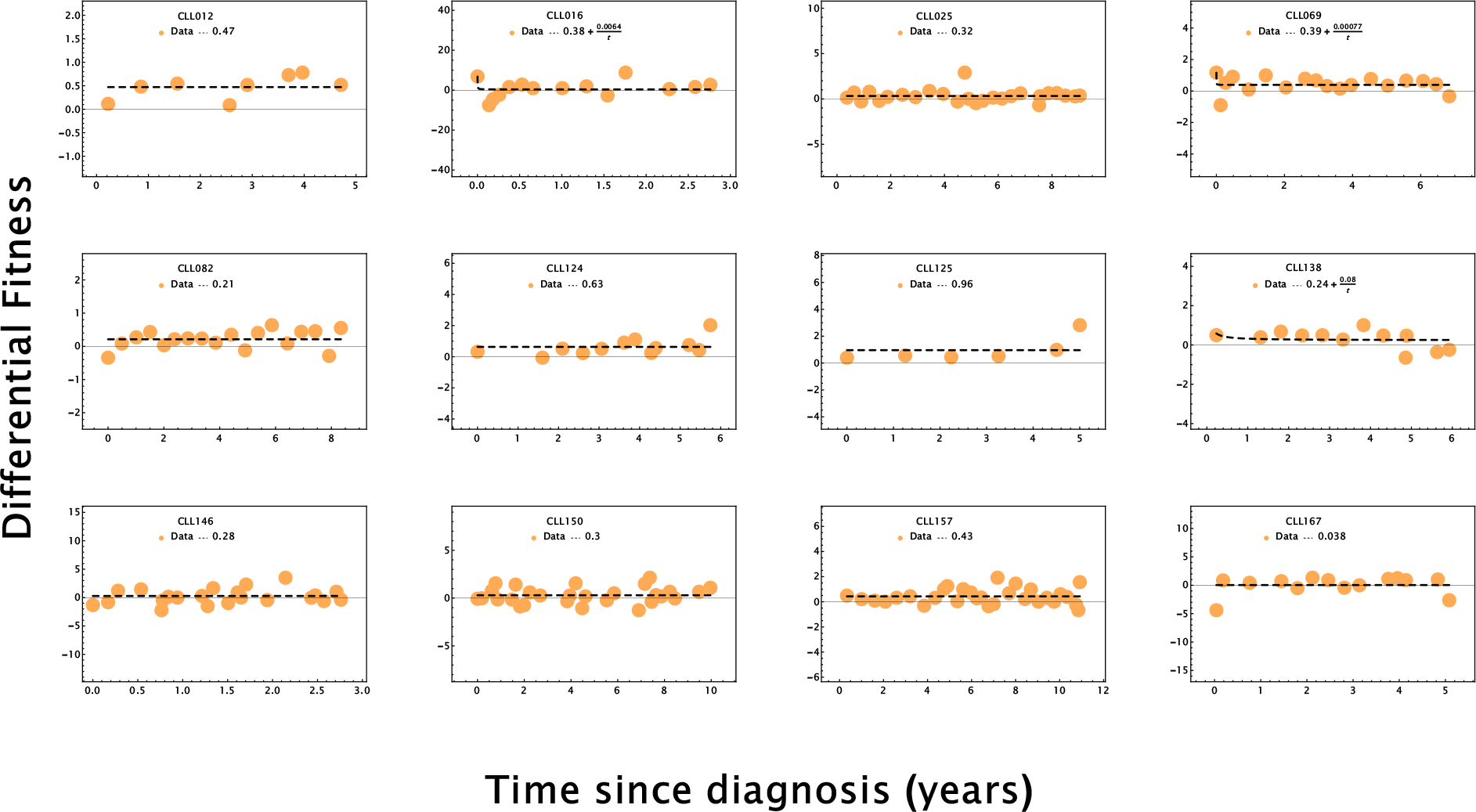
CCL patients with exponential growth pattern. The fitting is based on all available clinical observations.

### First 5−points fitting

**Figure S8.**
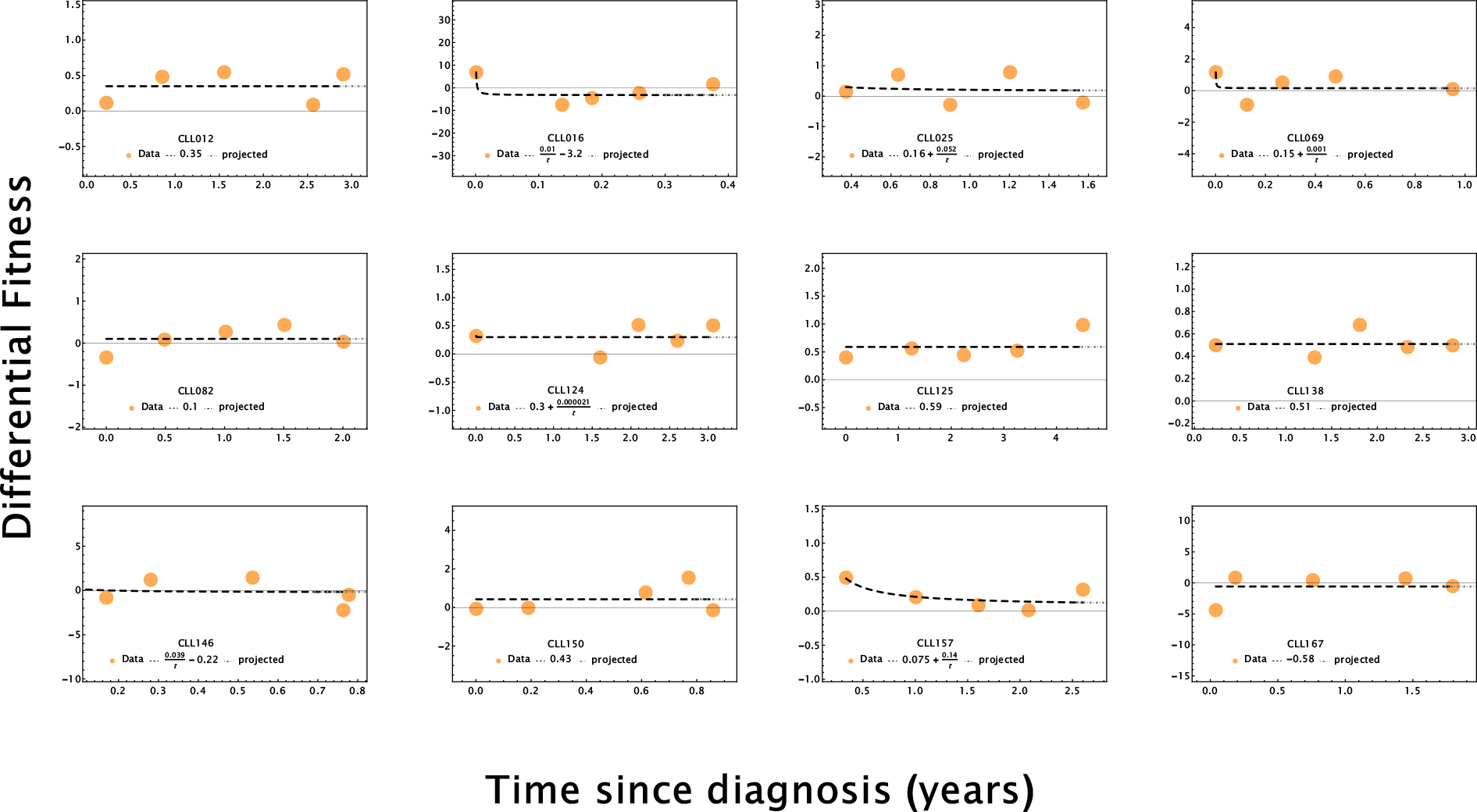
CCL patients with exponential growth pattern. The fitting is based on the first 5 clinical observations.

### First 7−points fitting

**Figure S9.**
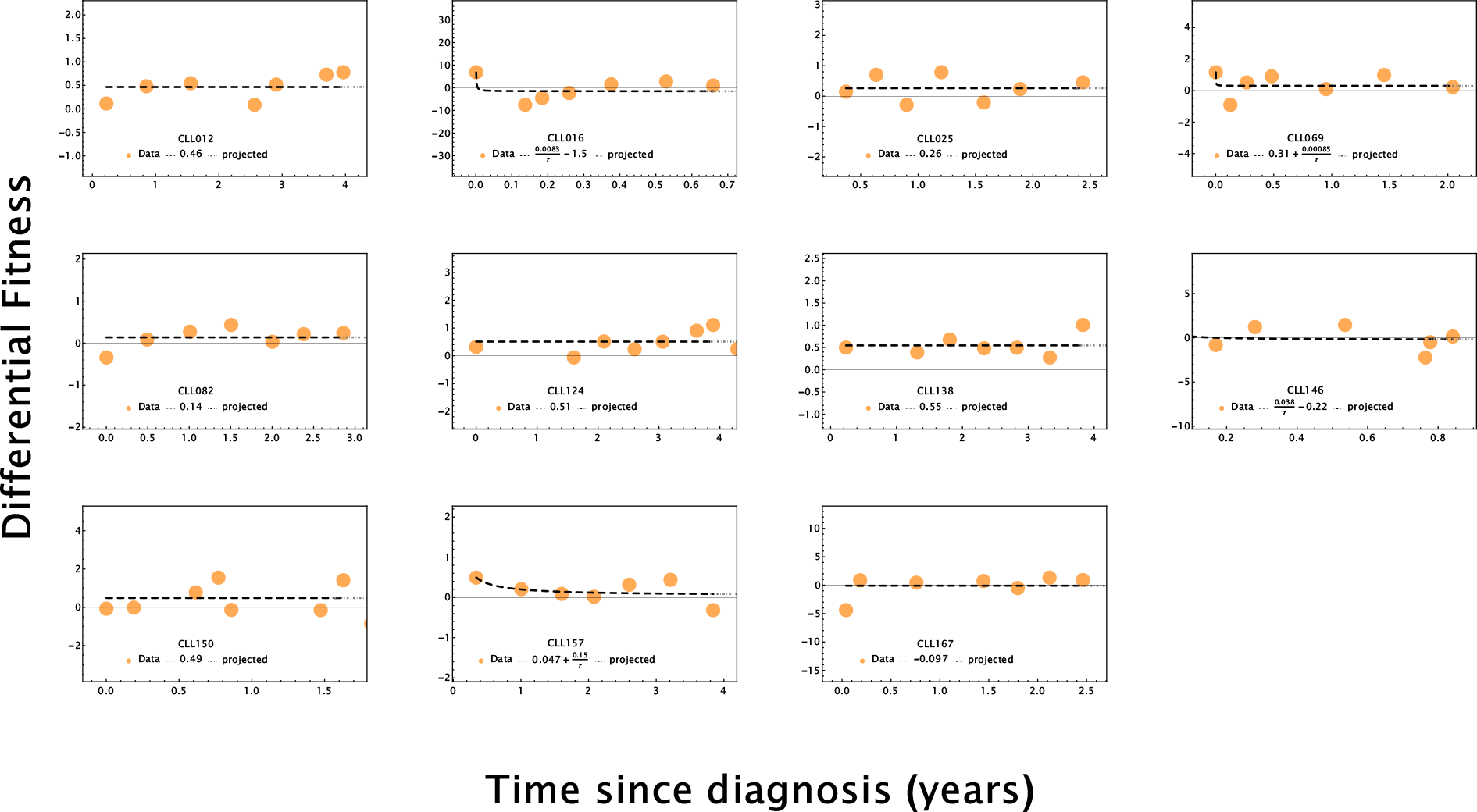
CCL patients with exponential growth pattern. The fitting is based on the first 7 clinical observations.

### First 10−points fitting

**Figure S10.**
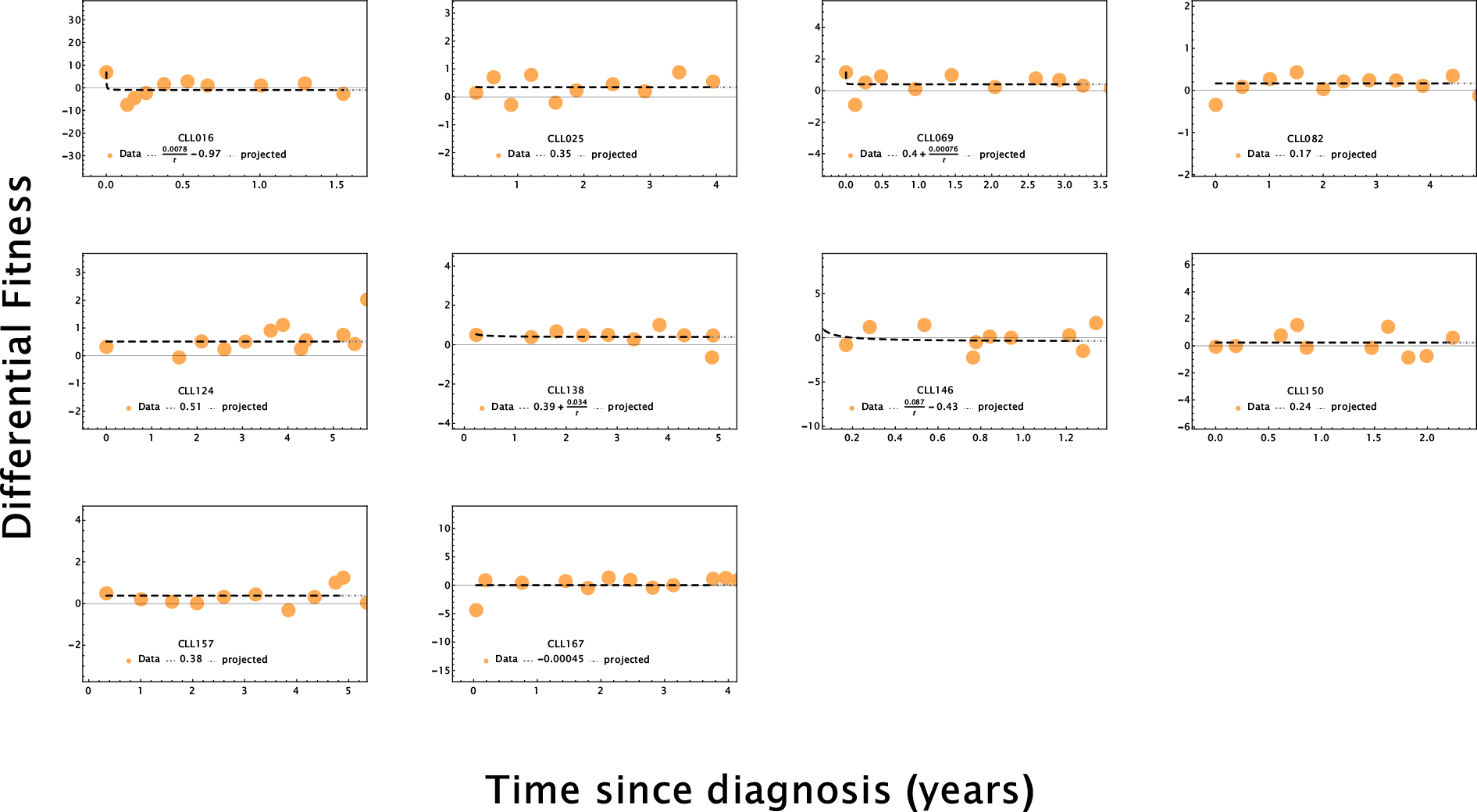
CCL patients with exponential growth pattern. The fitting is based on the first 10 clinical observations.

### CLL indeterminate growth pattern

**All points fitting**

**Figure S11.**
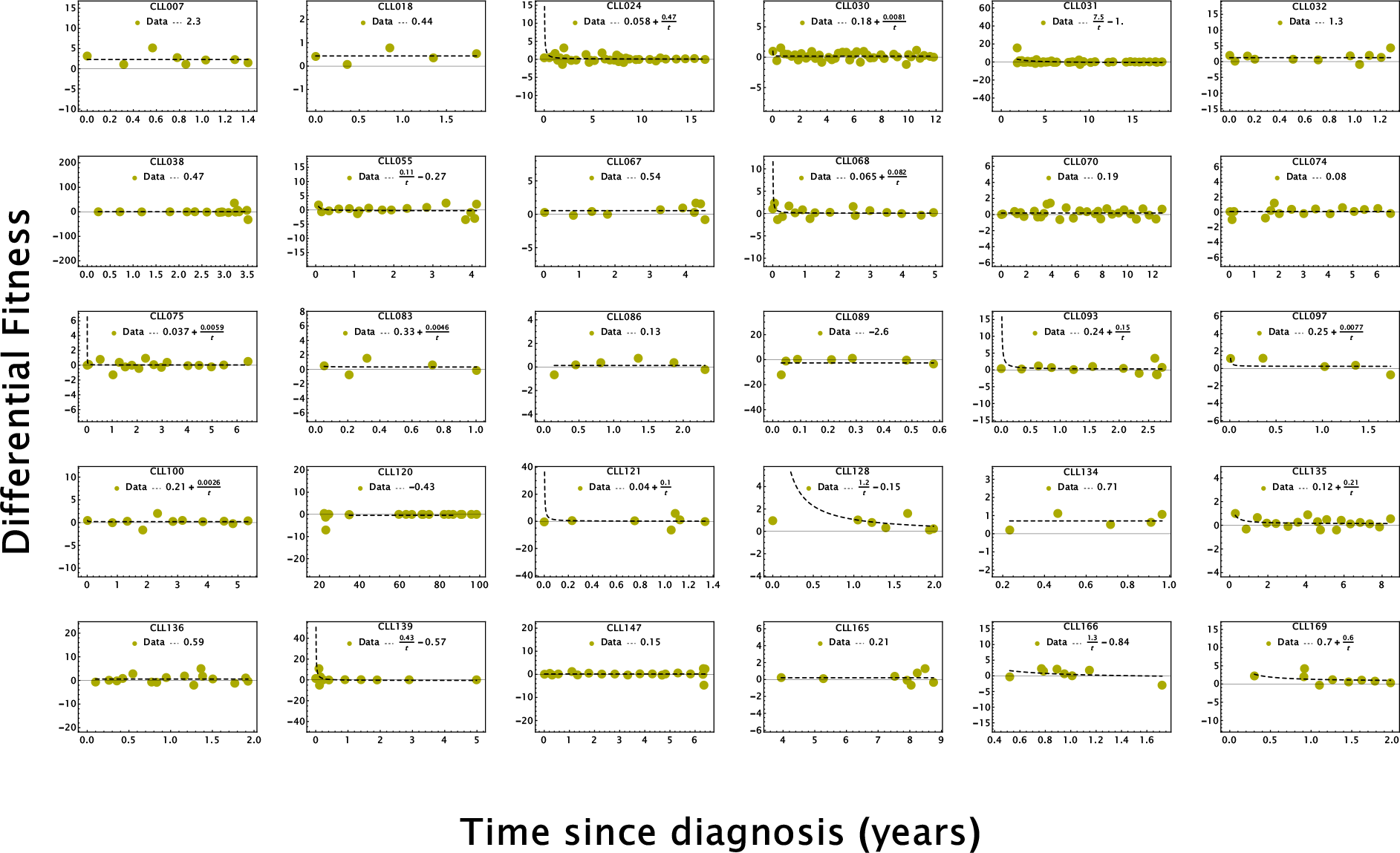
CCL patients with indeterminate growth pattern. The fitting is based on all available clinical observations.

### First 5–points fitting

**Figure S12.**
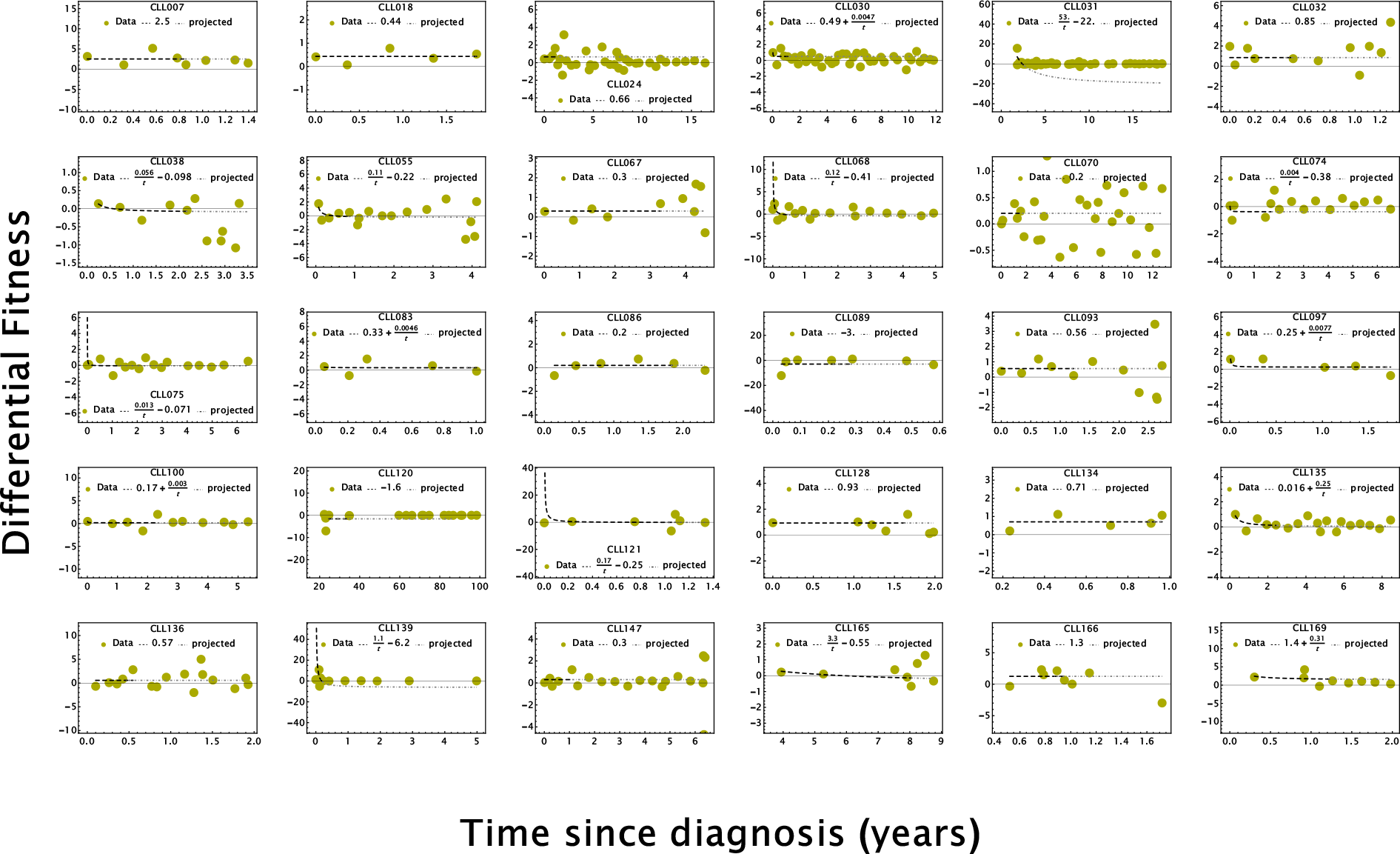
CCL patients with indeterminate growth pattern. The fitting is based on the first 5 clinical observations.

### First 7–points fitting

**Figure S13.**
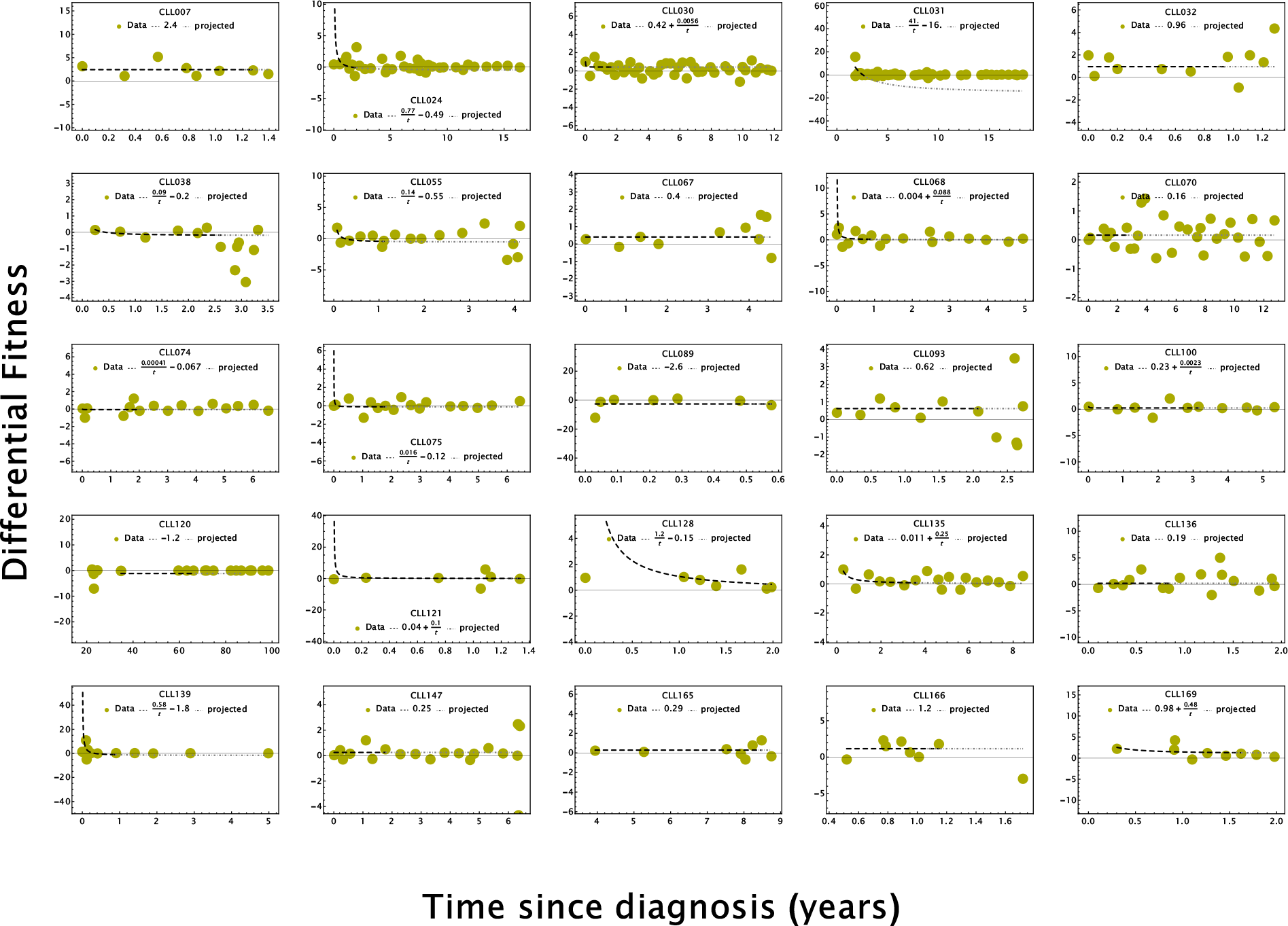
CCL patients with indeterminate growth pattern. The fitting is based on the first 7 clinical observations.

### First 10–points fitting

**Figure S14.**
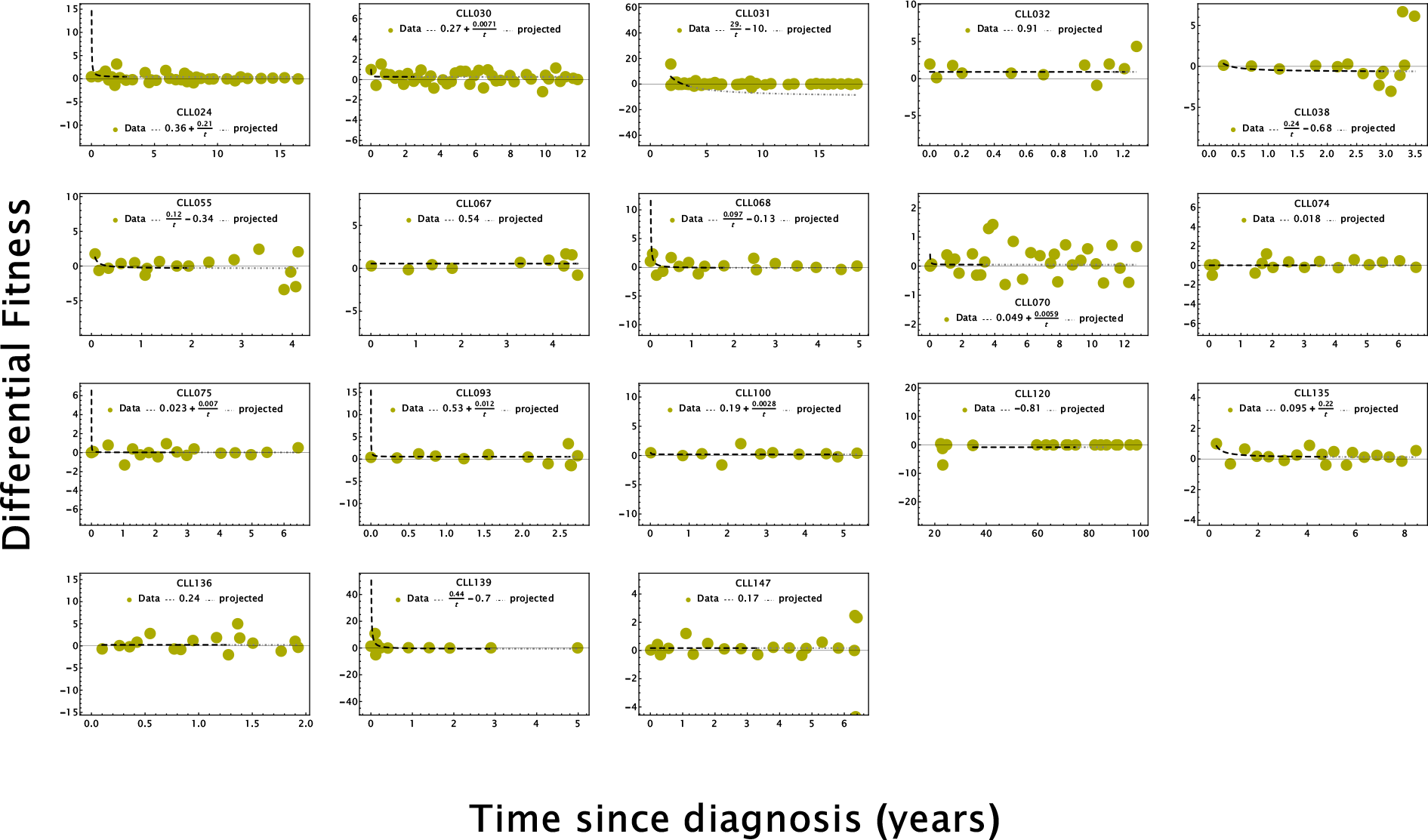
CCL patients with indeterminate growth pattern. The fitting is based on the first 10 clinical observations.

**Figure S15.**
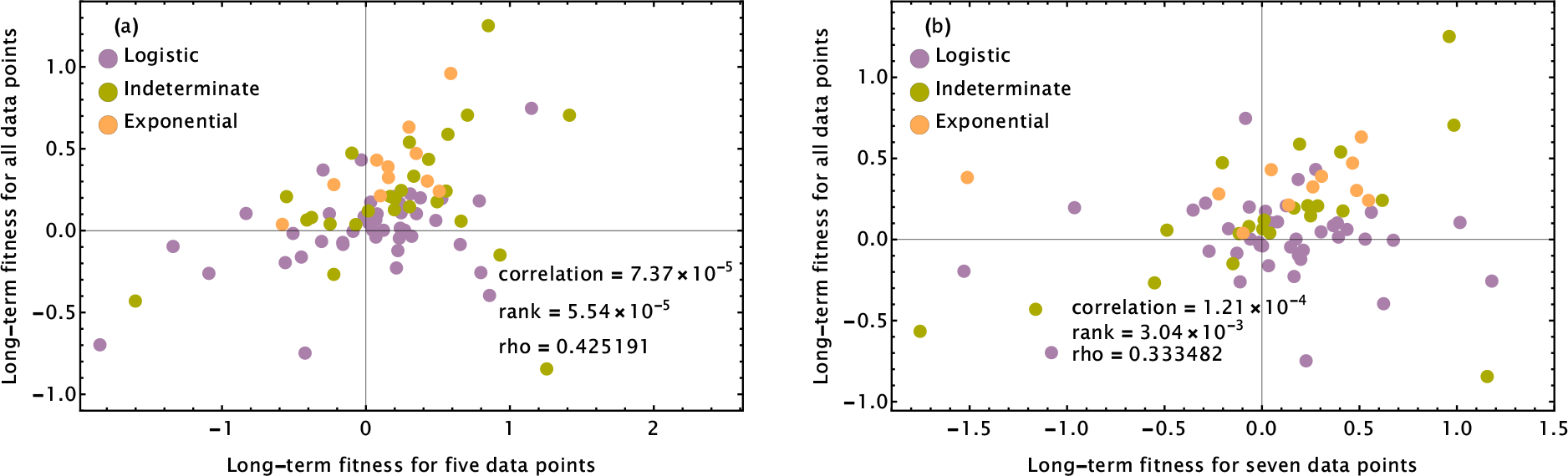
Pearson Correlation coefficient and Spearman Rho values for the three growth patterns plotted for the first 5 and 7 readings vs all readings of CLL patients respectively.

**Figure S16.**
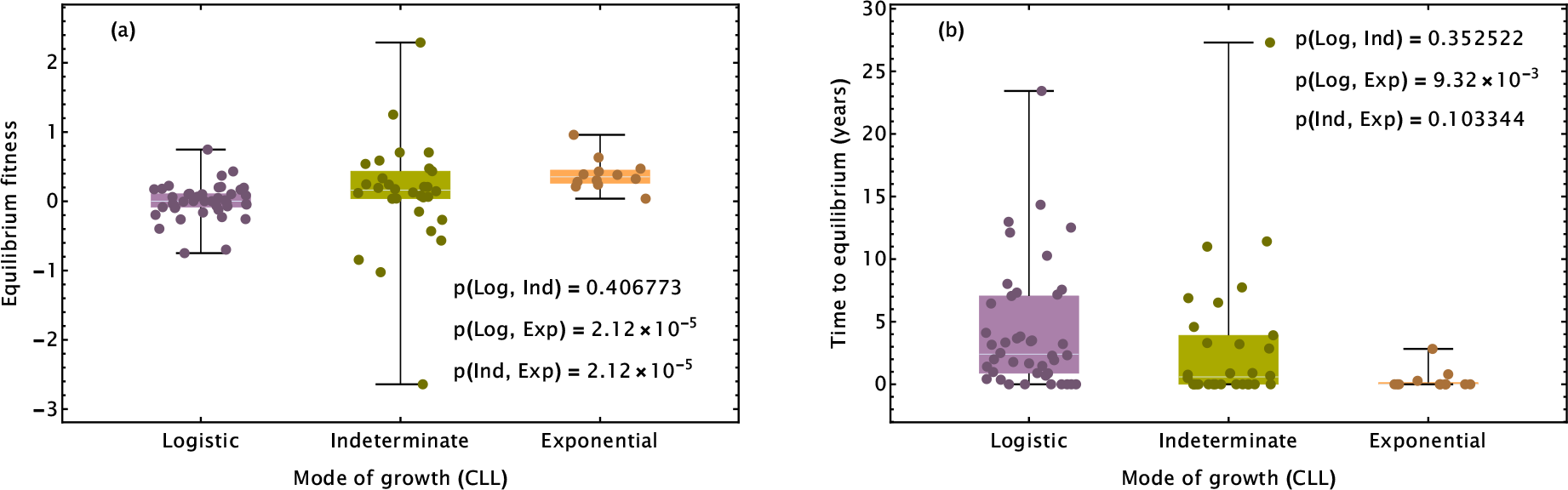
Box Whisker chart for the logistic, indeterminate, and exponential growth pattern.

